# Instability of Alpha Oscillatory States in Autism and Familial Liability: Evidence from Burst-Resolved High-Density Electroencephalography (EEG)

**DOI:** 10.64898/2026.04.03.716324

**Authors:** Theo Vanneau, Chloe Brittenham, Megan Darrell, Michael Quiquempoix, John J. Foxe, Sophie Molholm

**Affiliations:** The Cognitive Neurophysiology Laboratory, Departments of Pediatrics & Neuroscience, Albert Einstein College of Medicine, Bronx, New York 10461, USA; Institut de Recherche Biomédicale des Armées (IRBA), 91223 Brétigny sur Orge, France; URP 7330 VIFASOM, Université Paris Cité, Hôtel Dieu, Paris, France; The Frederick J. and Marion A. Schindler Cognitive Neurophysiology Laboratory, The Ernest J. Del Monte Institute for Neuroscience, Department of Neuroscience, University of Rochester School of Medicine and Dentistry, Rochester, New York 14642, USA

## Abstract

Atypical sensory experiences are highly prevalent in autistic children and include both hyper- and hypo-responsivity, often accompanied by sensory overload. Alpha oscillations (7–13 Hz), which dynamically regulate cortical excitability, represent a plausible neural mechanism underlying these phenomena: reduced alpha activity is associated with enhanced sensory responsiveness, whereas increased alpha supports suppression of external input. Although decreased alpha power has been repeatedly reported in autism, it remains unclear whether this reduction reflects lower oscillatory amplitude or reduced temporal stability of alpha rhythms, two mechanisms with distinct neurophysiological implications. To better characterize alpha activity in autism, we examined resting-state alpha dynamics in non-autistic children (NA; n = 39), autistic children (AU; n = 52), and siblings of autistic children (SIB; n = 26), aged 8–14 years. We combined traditional broadband measures of relative alpha power, parametric separation of periodic and aperiodic activity, and single-event analyses that quantify the temporal structure of alpha oscillations. Both broadband relative alpha power and periodic alpha power were reduced in autism over parietal regions, replicating prior findings. Importantly, ordinal analyses revealed an intermediate profile in siblings, supporting a liability-related gradient of alpha alterations. However, single-event analyses demonstrated that the average amplitude of individual alpha bursts did not differ between groups. Instead, autistic children showed significantly shorter alpha burst duration and reduced alpha abundance (i.e., proportion of time occupied by rhythmic alpha episodes), with siblings again exhibiting intermediate values. Linear regression analyses confirmed that reductions in relative and periodic alpha power were primarily driven by decreased alpha abundance rather than diminished burst amplitude. These findings indicate that altered alpha activity in autism reflects reduced temporal stability and density of alpha events rather than weaker oscillatory amplitude per se. Reduced persistence of alpha rhythms may therefore represent a neural marker of altered cortical excitability and sensory regulation in autism.

**Lay summary:** Autistic children often experience the world differently at the sensory level, including being more easily overwhelmed by sounds, lights, or other stimuli. In this study, we looked at a type of brain activity called alpha rhythms, which help regulate how strongly the brain responds to incoming information. We found that, in autistic children, these alpha rhythms were not weaker when they occurred, but they lasted for a shorter time and happened less often. Siblings of autistic children showed an intermediate pattern. These results suggest that sensory differences in autism may be linked to less stable brain rhythms that normally help control sensory input.

**Graphical Abstract:** 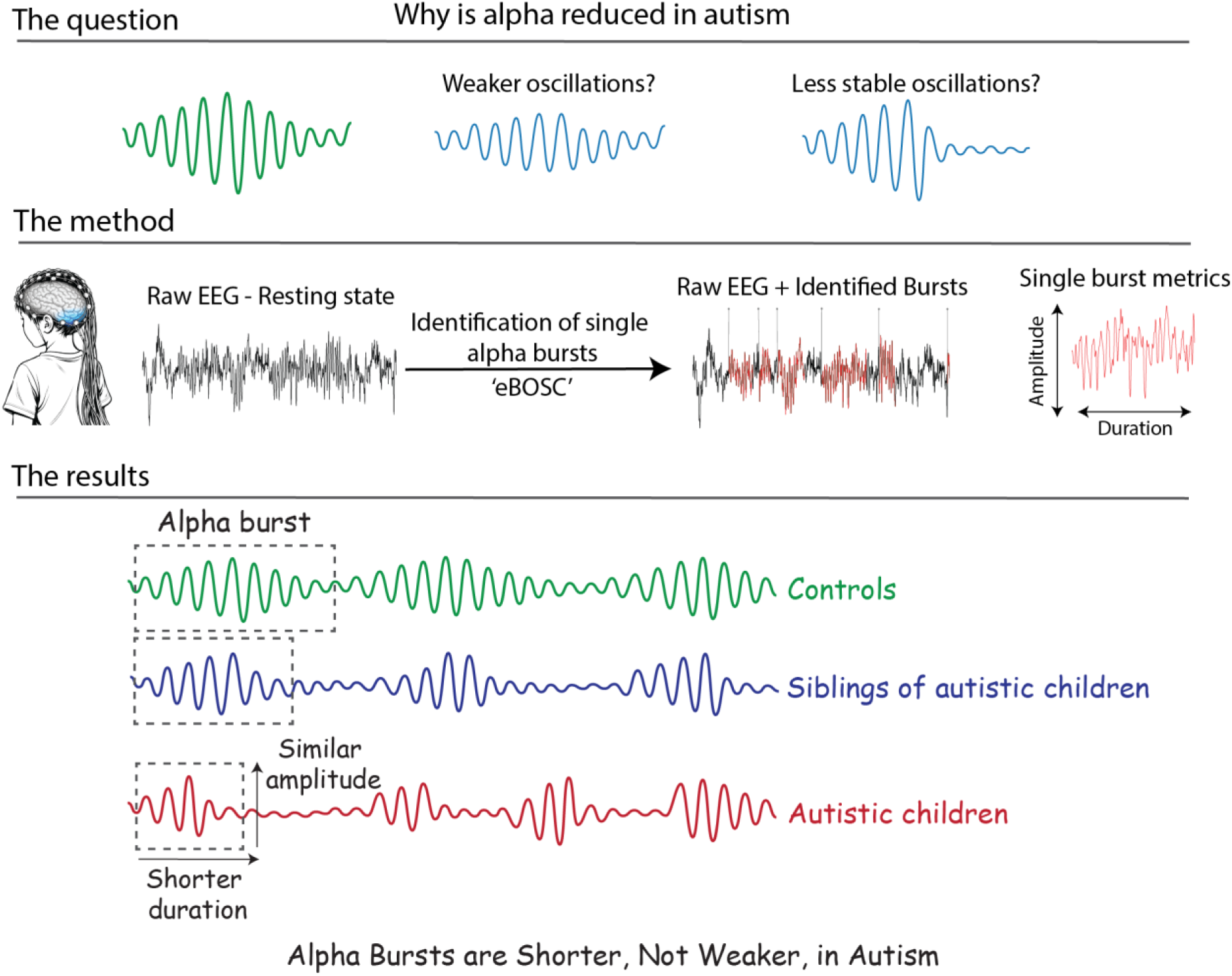

## Introduction

Autism spectrum disorder (ASD) is characterized not only by differences in social communication and restricted behavior, but also by pervasive atypical sensory experiences (American Psychiatric, 2022; Robertson & Baron-Cohen, 2017). Many autistic individuals report heightened sensitivity or feeling overwhelmed in everyday environments (e.g., classrooms or supermarkets), suggesting difficulties prioritizing processing of meaningful over distracting incoming sensory information (Belmonte, 2017; MacLennan et al., 2022; Taels et al., 2023).

Effective regulation of sensory processing depends on the dynamic modulation of cortical excitability, a process strongly shaped by alpha-band neuro-oscillatory activity (∼7-13 Hz). Converging evidence from human scalp and intracranial EEG recordings suggests that alpha activity reflects rhythmic processes that modulate cortical and thalamocortical excitability (Haegens et al., 2011; Mathewson et al., 2011), with alpha rhythms widely considered a top-down cortical sensory gating mechanism (Foxe et al., 1998; Iemi et al., 2022). Increases in alpha power are associated with reduced excitability and consequent suppression of sensory processing, whereas decreases in alpha power reflect heightened excitability and facilitate perception (Foxe & Snyder, 2011; Kelly et al., 2009; Sauseng et al., 2005; Worden et al., 2000). Importantly, alpha activity does not typically manifest as a sustained stationary oscillation. Rather, it occurs as transient, intermittent bursts, brief episodes of rhythmic synchronization that emerge and dissipate over time (Donoghue et al., 2022; Hanslmayr et al., 2011; Jones, 2016; Kosciessa et al., 2020; Steriade et al., 1990). These bursts are thought to reflect periods during which neural populations transiently synchronize at alpha frequency, producing rhythmic fluctuations in cortical excitability (Klimesch et al., 2007). This temporal organization has important implications: alpha is better conceptualized not as a continuous background rhythm, but as a sequence of discrete oscillatory events defined by their amplitude, duration, frequency, and timing (van Ede et al., 2018; Vidaurre et al., 2016).

A burst-based framework has been successfully applied in other frequency bands, such as beta and gamma, where single-trial analyses during memory tasks have demonstrated that oscillatory activity consists of transient burst events characterized by their rate, duration, and timing (Feingold et al., 2015; Lundqvist et al., 2018; Lundqvist et al., 2024; Sherman et al., 2016). These features index separable aspects of underlying neural physiology rather than merely contributing to average power.

Within the alpha band, burst amplitude may reflect the strength or spatial extent of inhibitory engagement (that is, the size of the recruited neural ensemble) whereas burst duration may indicate how long cortical excitability remains biased toward inhibition (Klimesch et al., 2007). Burst occurrence and abundance may reflect the probability and stability of entering an alpha oscillatory state. Because these parameters capture partially distinct neurophysiological processes, examining them independently provides a more mechanistic characterization of alpha-mediated sensory regulation.

Importantly, the expression of these oscillatory states can be studied both during task engagement and during spontaneous brain activity. Resting-state recordings provide a complementary perspective on alpha dynamics. During tasks, alpha modulation reflects rapid adjustments of cortical excitability driven by external demands and task goals (Vanneau et al., 2025). In contrast, resting-state activity captures the spontaneous organization of neural dynamics in the absence of explicit behavioral constraints (Klimesch, 1999). Resting alpha is therefore thought to reflect baseline fluctuations in cortical excitability and inhibitory regulation across large-scale cortical networks (Jensen & Mazaheri, 2010; Mathewson et al., 2011; Sadaghiani & Kleinschmidt, 2016). Examining burst-resolved alpha activity at rest makes it possible to characterize how frequently and how stably the brain engages in these inhibitory oscillatory states, independently of task-driven modulation.

Given the central role of alpha oscillations in regulating cortical excitability and sensory gating, alterations in alpha dynamics may contribute to the perceptual and attentional differences frequently reported in autism. Several domains in which autistic individuals show perceptual and attentional differences map closely onto known functions of alpha oscillations. These include altered attentional filtering (Dawson, Webb, et al., 2004; Keehn et al., 2013; Murphy et al., 2014) and reduced perceptual switching in multistable tasks such as binocular rivalry (Robertson et al., 2016). Despite this conceptual convergence, findings on resting-state alpha power in autism remain inconsistent. A recent meta-analysis of 41 studies reported no group differences in absolute alpha power, however they report a reduction in relative alpha power (Neo et al., 2023), a pattern echoed by several narrative reviews (Billeci et al., 2013; Newson & Thiagarajan, 2018; Wang et al., 2013).

However, conventional power estimates collapse activity across time and therefore cannot determine whether reduced relative alpha power reflects weaker oscillatory amplitude or reduced temporal engagement in an oscillatory state. These alternatives imply distinct neurophysiological mechanisms. A reduction in oscillatory amplitude would suggest diminished recruitment or synchronization of inhibitory neural populations, potentially reflecting weaker inhibitory engagement. In contrast, reduced time spent in rhythmic alpha states would indicate that inhibitory oscillatory periods occur less frequently or are less sustained over time, resulting in shorter or less stable inhibitory windows (Hanslmayr et al., 2011). Distinguishing between these possibilities is essential for clarifying the underlying neural mechanism and for informing targeted neuromodulatory approaches, such as neurofeedback or transcranial magnetic stimulation.

To date, no study has systematically examined burst-resolved alpha features in autism at rest. To address this gap, alpha activity was first characterized using conventional spectral approaches by estimating time-averaged alpha power, including both broadband and periodic components after separating periodic from aperiodic activity (Donoghue et al., 2020). While such measures provide a useful summary of oscillatory engagement, they collapse activity across time and therefore cannot reveal which underlying temporal features contribute to potential group differences. To resolve this, an episode-based approach was then applied (Kosciessa et al., 2020; Whitten et al., 2011) to identify discrete alpha bursts and quantify their amplitude, frequency, duration, occurrence, and abundance. By combining conventional spectral estimates with burst-resolved metrics, this framework makes it possible to determine which specific temporal features of alpha activity account for differences observed in average alpha power. Treating resting-state EEG as a continuous time series, burst-resolved parameters were extracted during a 1-minute eyes-open resting-state recording, allowing a detailed characterization of intrinsic alpha dynamics in autism.

We further examined typically developing individuals with autistic siblings (SIBs), who share familial risk but not clinical diagnosis, to test whether alpha dynamics exhibit intermediate profiles consistent with an endophenotype. Notably, resting-state alpha power has been linked to autistic traits related to behavioral rigidity in typically developing adults (Carter Leno et al., 2018), suggesting that alpha alterations may extend beyond diagnosed individuals and reflect broader familial liability. Including siblings allows us to distinguish disorder-specific alterations from neural traits associated with genetic risk.

Based on frameworks positioning alpha as a cortical sensory gating mechanism (Dockree et al., 2007; Foxe & Snyder, 2011; Klimesch et al., 2007; Van Diepen et al., 2019), we hypothesized that autistic children would show reduced alpha engagement at rest. Specifically, we predicted reduced time-averaged alpha power (both broadband and periodic components). When decomposing activity into burst-resolved metrics, however, we did not make a directional prediction regarding which specific burst features would account for this reduction. Decreased alpha power could arise from weaker burst amplitude, fewer burst events, shorter burst durations, or a combination of these factors, each reflecting a distinct alteration in inhibitory dynamics. For example, reduced burst amplitude could indicate weaker inhibitory recruitment, whereas reduced burst occurrence or duration would suggest that inhibitory oscillatory states are entered less frequently or sustained for shorter periods. Either pattern would be consistent with clinical reports of sensory hyper-responsivity and increased distractibility in autism (Dawson, Toth, et al., 2004; Keehn et al., 2013; Keehn et al., 2016; Keehn et al., 2017). By resolving alpha activity into its burst components, this approach allows identification of the specific temporal features underlying potential group differences, thereby providing mechanistic insight into the regulation of cortical excitability in autism.

## Methods

### Participants

The study initially included 41 non-autistic (NA), 54 autistic (AU), and 28 sibling (SIB) participants, all between 8 and 14 years of age. After excluding participants based on EEG quality, the final analysis was conducted on 39 NA, 52 AU and 26 SIB (see Table 1 for participant characteristics). To be included in the AU group, participants had to meet diagnostic criteria for AU on the basis of the following measures: *1*) autism diagnostic observation schedule 2 (ADOS-2) (Lord et al., 1994); *2*) diagnostic criteria for autistic disorder from the *Diagnostic and Statistical Manual of Mental Disorders* (DSM-5 (American Psychiatric, 2022)); *3*) clinical impression of a licensed clinician with extensive experience in diagnosis of children with ASD. Due to precautions during the COVID-19 pandemic, a subset of AU participants (n=9) was not able to complete the ADOS-2 (Lord et al., 1994), as masking requirements impacted administration. These participants instead underwent the Childhood Autism Rating Scale 2 (CARS-2) and Autism Diagnostic Interview-Revised (ADI-R) (Rutter et al., 2003) for confirmation of diagnosis. Participants in the NA group met the following inclusion criteria: no history of neurological, developmental, or psychiatric disorders, no first-degree relatives diagnosed with AU, and enrollment in age-appropriate grade in school. SIB group participants met the same criteria as the NA group, except that they had a sibling diagnosed with AU. Exclusion criteria for all groups included: (1) a known genetic syndrome associated with an IDD (including syndromic forms of AU), (2) a history of or current use of medication for seizures in the past 2 years, (3) significant physical limitations (e.g., vision or hearing impairments, as screened over the phone and on the day of testing), (4) premature birth (<35 weeks), or (5) a Full Scale IQ (FS-IQ) of less than 80. All procedures were approved by the Institutional Review Board of the Albert Einstein College of Medicine and adhered to the ethical standards outlined in the Declaration of Helsinki. All participants assented to the procedures, and their parents or guardian signed an informed consent approved by the Institutional Review Board of the Albert Einstein College of Medicine. Participants received nominal recompense for their participation (at $15 per hour).

**Table 1.** Demographic characteristics. NA: non-autistic control group; AU: autistic group; SIB: unaffected siblings of autistic individuals; FSIQ: Full-scale Intelligent coefficient; SRS-2: Social Responsiveness Scale, Second Edition, total score; ADOS-2: Autism Diagnostic Observation Schedule 2, total score.

| | NA (n=39) | AU (n=52) | SIB (n=26) | Statistic | p-value | $\eta^2$ |
| --- | --- | --- | --- | --- | --- | --- |
| **Sex**<br>F\|M | 18 \| 21 | 11 \| 41 | 15 \| 11 | $\chi^2 = 13.85$ | 0.007 | - |
| **Age**<br>Mean±Std<br>(range) | 10.50 ± 1.81<br>(8-14) | 10.73 ± 1.53<br>(8-14) | 10.56 ± 1.44<br>(8-14) | F=0.24 | 0.78 | 0.004 |
| **FSIQ**<br>Mean±Std | 103.63 ± 16.42 | 94.84 ± 16.14 | 103.0 ± 15.59 | F=3.92 | 0.02 | 0.067 |
| **SRS2** | 47.03 ± 6.57 | 72.18 ± 11.90 | 47.65 ± 7.70 | F = 93.01 | <0.001 | 0.63 |
| **ADOS** | - | 7.89 ± 2.1 | - | - | - | - |

### Experimental procedure

Participants were seated in a chair in an electrically shielded room (International Acoustics Company, Bronx, New York), 70 cm away from the visual display (Dell UltraSharp 1704FPT). 1-min eyes-open resting-state was recorded while the participant was instructed to watch the screen (gray screen with a fixation cross).

### EEG recordings & preprocessing

EEG data were recorded at a sampling rate of 512Hz using 64 channels BioSemi Active II system (using the CMS/DRL referencing system) with an anti-aliasing filter (−3 dB at 3.6 kHz). Analyses were conducted in python (3.11) using MNE (originally for Minimum Norm Estimation)(Gramfort et al., 2013) and custom scripts available at https://github.com/tvanneau/SFARI-RS-Alpha. Bad channel detection was performed using the function NoisyChannels (with RANSAC) from the pyprep toolbox (Bigdely-Shamlo et al., 2015). If more than 15% of the channels were detected as bad, the participants were rejected (2 NA; 2 AU and 2 SIB). Bad channels were interpolated using spline interpolation (Perrin et al., 1989). EEG was filtered using a FIR band-pass filter (0.01-40Hz), and Independent Component Analysis (ICA) on 1Hz high-pass EEG was used to identify and manually reject eye-related components (blinks/saccades).

### Broadband oscillatory metrics

Broadband spectral metrics were estimated for each channel and each participant from resting-state EEG using Welch’s method. Power spectral density (PSD) was computed using sliding windows of 2 seconds with 50% overlap and then interpolated onto a common frequency grid (1–35 Hz). For canonical frequency bands (theta: 3–7 Hz; alpha: 7–13 Hz; low beta: 13–20 Hz; high beta: 20–35 Hz), absolute and relative power were computed. Absolute power was obtained by integrating the power spectral density (PSD) within each band using the trapezoidal rule (area under the curve). Relative power was defined as the absolute band power divided by the total power across the 3–35 Hz range. Within the alpha band, additional metrics were derived: the peak alpha frequency (PAF), defined as the frequency exhibiting maximal power between 7 and 13 Hz along with its corresponding amplitude, and the individual alpha frequency (IAF), computed as the spectral barycenter of the alpha band, corresponding to the power-weighted mean frequency.

### Periodic and aperiodic spectral metrics

To dissociate oscillatory (periodic) activity from the arrhythmic background component, the same PSD was parameterized using the specparam algorithm (Donoghue et al., 2020) with a *knee* model for the aperiodic component. From the aperiodic fit, we extracted: the offset, the exponent and the knee parameter. From the aperiodic fit, three parameters were extracted: the offset, the exponent, and the knee. The periodic component was then analyzed to identify spectral peaks whose center frequency fell within the alpha range (7–13 Hz). Because removal of the aperiodic component can reveal multiple oscillatory peaks within this frequency range, all detected peaks were retained for further characterization. For these peaks, the following metrics were derived: the periodic individual alpha frequency (IAF; calculated after removal of the aperiodic component), the power above the aperiodic fit (PW), the bandwidth of each peak (BW), the number of detected alpha peaks, and the summed power of all periodic alpha peaks. All metrics were computed at the single-channel level and subsequently used for topographical modeling and cluster-based statistical analyses.

### Single-burst metrics

Alpha events were identified using the extended Better OSCillation detection method (eBOSC)(Caplan et al., 2001; Kosciessa et al., 2020; Whitten et al., 2011). First, the continuous EEG signal was transformed into the time–frequency domain using wavelets. The resulting power spectrum was then used to estimate the background 1/f activity after removal of the alpha peak. An amplitude threshold was subsequently derived from this background estimate and set to the 95th percentile of the modeled background distribution. Time–frequency points whose power exceeded this threshold were classified as rhythmically active (assigned a value of “1”), whereas remaining points were labeled as non-rhythmic (“0”). Next, eBOSC applies a duration criterion to ensure that detected rhythmic episodes reflect sustained oscillatory activity rather than brief high-amplitude transients. Specifically, the amplitude threshold was required to be exceeded for at least two consecutive cycles of the local alpha frequency (200 ms at 10 Hz), preventing transient fluctuations from being misclassified as oscillatory episodes. Time points meeting both the amplitude and duration criteria were merged into discrete alpha bursts. For each detected burst, the onset and offset times, mean frequency, mean power, and duration were extracted. These burst-resolved metrics enable characterization of alpha activity in terms of oscillatory amplitude, temporal persistence, and occurrence rate, independently of the aperiodic 1/f background component.

### Statistical analysis

To account for the intermediate liability profile of the SIB group, linear mixed-effects models (LMMs) were implemented using two complementary approaches. In the first approach, group membership was modeled as an ordinal factor reflecting a dimensional trend across groups (AU = −1, SIB = 0, NA = +1). This coding enabled estimation of a linear slope describing the monotonic change in each alpha parameter across the three groups and allowed testing whether this slope differed from zero. In the second, more classical approach, group was treated as a categorical factor with AU as the reference level, enabling pairwise comparisons between groups. For both modeling strategies, analyses were performed at the single-channel level. Channels were assigned to predefined anatomical clusters corresponding to frontal, centroparietal, and occipital regions, each subdivided by hemisphere (left/right). Single-channel observations within these clusters were treated as repeated measures within participants, allowing the model to capture spatial variability while accounting for within-subject dependence. Models were fitted using restricted maximum likelihood with the Limited-memory Broyden–Fletcher–Goldfarb–Shanno (L-BFGS) optimizer. Fixed effects included group (or the ordinal trend), region, hemisphere, and their interactions with the group term. Age and sex were included as covariates in all models. A random intercept was specified for each participant. This modeling framework was applied to alpha metrics derived from Welch spectral power estimates, specparam-based periodic activity, and eBOSC burst parameters. Additional models were run including Full-Scale IQ (FSIQ) and Social Responsiveness Scale scores (SRS-2) as covariates. For all analyses, regression coefficients of significant fixed effects are reported together with their associated p-values (α = 0.05). Finally, to determine whether reductions in relative alpha power were better explained by changes in burst occurrence or burst magnitude, multiple linear regression analyses were conducted in which relative alpha power served as the dependent variable and burst abundance and burst amplitude were entered simultaneously as predictors. The contribution of each predictor was evaluated using standardized regression coefficients and by examining relationships among residualized variables.

## Results

Resting-state alpha activity was first examined using conventional approaches that quantify oscillatory power averaged across time. The analysis tested whether autistic (AU) participants showed the commonly reported reduction in alpha power relative to non-autistic (NA) participants, using both broadband relative alpha power and periodic alpha power after separation from aperiodic (1/f) activity. In addition, unaffected siblings (SIB) were evaluated for intermediate values, consistent with liability or endophenotype models.

To address these questions, a hierarchical statistical strategy was adopted. Diagnostic status was first modeled as an ordinal factor (AU → SIB → NA) to test graded progression across groups. A significant slope in this model indicates that siblings fall between autistic and non-autistic participants, consistent with a familial liability continuum. Only when a significant ordinal effect was observed was a categorical model subsequently implemented, treating group as a nominal factor. This enabled direct pairwise comparisons and quantification of the magnitude of separation between specific groups while limiting unnecessary multiple testing.

### Reduced parieto-occipital alpha activity in autism with an intermediate profile in siblings

#### Absolute alpha power

For absolute alpha power, the ordinal (slope) model revealed no significant overall linear trend across groups (Supplementary Fig. 1B; β = 0.04, SE = 0.03, p = .25). However, regional effects were robust. Across participants, alpha power showed the expected posterior dominance, with significantly higher alpha over the occipital region relative to parietal scalp (β = 0.33, SE < 0.01, p < .001; alpha expressed in log units), and significantly lower alpha over frontal regions (β = −0.086, SE < 0.01, p < .001). Critically, a significant Group slope × Region interaction (β = −0.026, SE < 0.01, p < .001) indicated that the linear increase across groups was significantly attenuated at frontal sites relative to centroparietal regions. Thus, siblings displayed an intermediate profile primarily over parieto-occipital areas.

Age was significantly associated with absolute alpha power (Supplementary Fig. 1C; β = −0.066, SE = 0.01, p < .001), indicating a developmental decrease in alpha amplitude. However, there was no significant Age × Group slope interaction, suggesting that group differences in spatial alpha organization were stable across age. Sex was not significantly associated absolute alpha power and there was no significant Sex x Group slope interaction (p_s_ > 0.5).

The categorical model yielded converging results. Although there was no main effect of group, a significant Group × Region interaction was observed. Examination of the model coefficients indicated that this interaction was driven by differences in the frontal versus centroparietal contrast. Specifically, the reduction of alpha power at frontal sites relative to centroparietal regions was stronger in siblings (β = −0.056, SE = 0.01, p = .001) and neurotypical participants (β = −0.050, SE = 0.01, p = .001) compared with autistic participants (reference group). This pattern indicates that the posterior–frontal alpha gradient was attenuated in autism, with siblings showing an intermediate spatial profile (Supplementary Fig. 1D).

#### Relative power

In contrast, relative alpha power exhibited a significant positive slope (β = 2.6%, SE = 0.01, p = .026; Fig. 1D), indicating increasing alpha from AU → SIB → NA. Importantly, this effect was topographically specific. A significant slope × Region interaction demonstrated that the ordinal increase was markedly reduced at frontal sites compared with centroparietal regions (β = –1.3%, SE < 0.01, p < .001). Thus, siblings were most clearly intermediate (to AU and NA) over parieto-occipital areas. Age was not associated with relative alpha power (ß < 0.01, SE < 0.01, p = .61). Using the categorical approach, autistic participants differed significantly from controls (β = 5.1%, SE = 0.02, p = .027), whereas siblings did not differ from autism (β = 2.9%, SE = 0.02, p = .29). Sex was not significantly associated with relative power.

**Figure 1.**
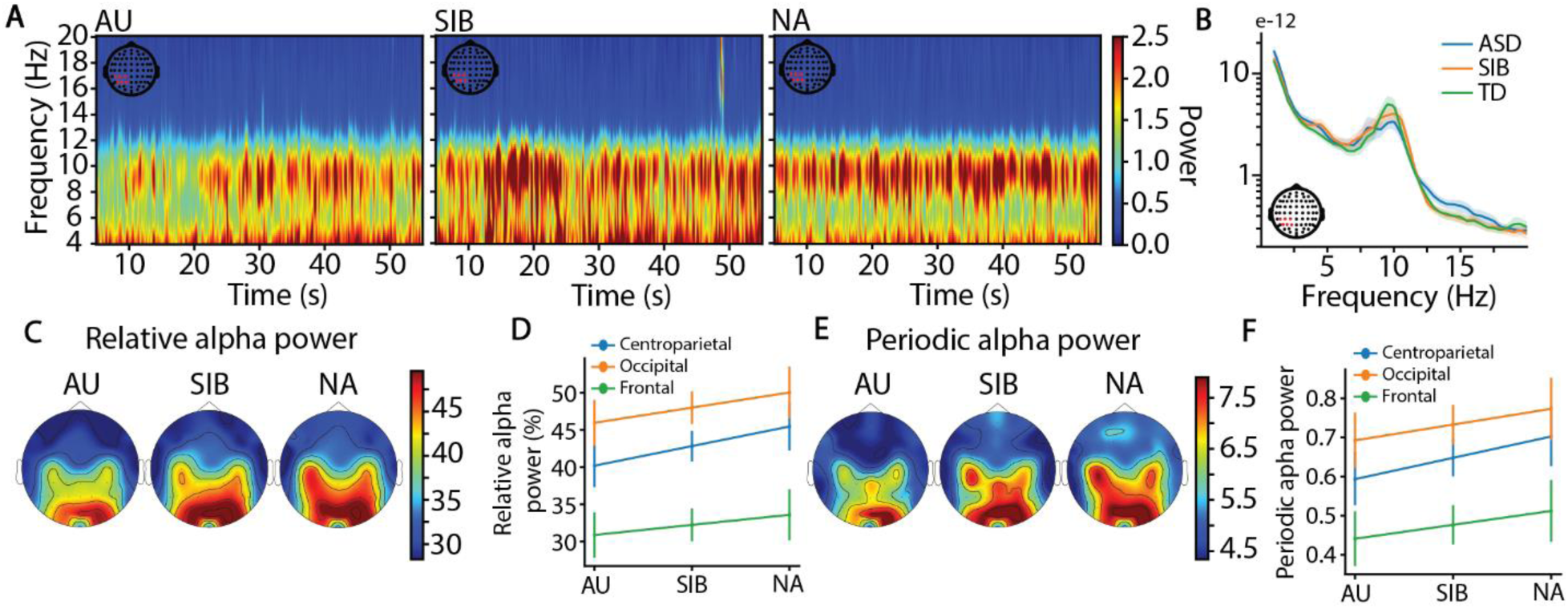
Reduced parieto-occipital alpha activity in autism with an intermediate profile in siblings. **(A)** Time–frequency representations averaged over the left centroparietal cluster for autistic (AU; left), sibling (SIB; middle), and non-autistic (NA; right) participants. (B) Power spectral density averaged across time for the same cluster, shown for AU (blue), SIB (orange), and NA (green). (C) Scalp topographies of relative alpha power for AU (left), SIB (middle), and NA (right). (D) Predicted values from the linear mixed model (LMM) illustrating the ordinal group trend in relative alpha power. (E–F) Same conventions as in panels C and D, but for periodic alpha power estimated after separation of periodic and aperiodic components using the specparam approach (Donoghue et al., 2020). The LMMs revealed a significant ordinal trend across groups (see Results).

#### Alpha frequency measures

Individual alpha frequency (IAF) did not vary across the ordinal axis (Supplementary Fig. 1E; ß = 0.027, SE = 0.23, p = 0.9) but there was an anterior-posterior gradient, with lower IAF for frontal channels (ß = −0.07, SE < 0.01, p < 0.001) and higher IAF for occipital channels (ß = 0.07, SE < 0.01, p < 0.001) relative to the parietal region, without significant interaction between group and region. In addition, the right hemisphere was associated with slower alpha burst frequency compared to the left hemisphere (β = −0.019 Hz, SE < 0.01, p = .006) and age was significantly associated with IAF (Supplementary figure. 1G; ß = 0.09, SE < 0.01, p < 0.001) while sex was not significantly associated with IAF.

#### Periodic versus aperiodic contributions

We next examined whether the observed liability pattern in alpha power could be accounted for by differences in aperiodic activity. Using parametric spectral decomposition (Donoghue et al., 2020), we separated periodic alpha peaks from the aperiodic background component of the spectrum.

#### Aperiodic Activity

For the aperiodic offset, we observed a robust anterior–posterior gradient across participants, with higher offset values over occipital regions (β = 0.50, SE = 0.02, p < .001) and lower values over frontal regions relative to the parietal region (β = −0.30, SE = 0.02, p < .001; Supplementary Fig. 1H). Importantly, no significant Group × Region interactions were found, indicating that this spatial gradient was comparable across groups. Age was not significantly associated with the aperiodic offset (β = −0.07, SE = 0.05, p = .159). A similar pattern emerged for the aperiodic exponent. We again observed a pronounced anterior–posterior gradient (Supplementary Fig. 1I), with higher exponent values in occipital regions (β = 0.20, SE = 0.01, p < .001) and lower values in frontal regions relative to parietal cortex (β = −0.30, SE = 0.01, p < .001). No group differences or interactions were detected, and age was not significantly related to the exponent (β = −0.01, SE = 0.03, p = .63). Sex was not significantly associated with either the aperiodic offset or the exponent. Together, these results indicate that aperiodic parameters follow a consistent posterior-dominant topography across groups and do not account for the observed liability-related effects in alpha power.

#### Periodic Alpha Power

In contrast, periodic alpha power reproduced the liability pattern. We observed a significant positive linear trend across groups (ASD → SIB → controls; β = 0.057, SE = 0.02, p = .04; Fig. 1E), indicating progressively higher periodic alpha power from autism to controls. Across participants, periodic alpha exhibited a robust anterior–posterior gradient, with greater power in occipital regions relative to parietal cortex (β = 0.085, SE < 0.01, p < .001) and reduced power in frontal regions (β = −0.17, SE < 0.01, p < .001). Critically, the liability effect was topographically specific. A significant Group trend × Region interaction (β = −0.019, SE < 0.01, p = .02) indicated that the linear increase across groups was significantly attenuated at frontal sites relative to centroparietal regions. Thus, siblings exhibited a clearly intermediate profile primarily over parieto-occipital areas. Age was not significantly associated with periodic alpha power (β < 0.001, SE = 0.01, p = .92). The categorical model yielded converging results: autistic participants differed from controls (β = 0.11, SE = 0.05, p = .04), whereas siblings did not differ significantly from the autism group (β = 0.05, p = .35), consistent with an intermediate familial profile. Sex was not significantly associated with periodic alpha power.

#### Periodic Alpha Frequency

The periodic IAF (IAF computed after removal of the aperiodic activity; p-IAF) followed the same pattern than the IAF, with the anterior-posterior gradient with lower p-IAF for frontal channels (Supplementary Fig. 1J; ß = −0.084, SE = 0.03, p < 0.01) and higher frequency for occipital channels (ß = 0.019, SE = 0.03, p = 0.57), without significant interaction between group and region, showing that this pattern is similar between groups. In addition, the right hemisphere was associated with slower alpha frequency compared to the left hemisphere (β = −0.023 Hz, SE < 0.01, p < .001). Age was significantly associated with p-IAF (Supplementary Fig. 1L; ß = 0.08, SE = 0.01, p < 0.001) without significant interaction with groups and sex was not significantly associated with p-IAF.

Across both broadband relative measures and parametrically isolated periodic components, alpha activity followed a consistent gradient: AU < SIB < TD, most prominently over parieto-occipital cortex. Siblings therefore occupied an intermediate position across independent analytical frameworks, supporting a quantitative liability account rather than a strictly categorical distinction between diagnostic groups.

### Temporal organization of alpha activity

Having established group differences in time-averaged alpha power, the analysis next focused on temporal dynamics. Conventional power estimates collapse across time and therefore cannot determine whether group differences reflect weaker oscillatory events or reduced time spent in an oscillatory state. To disentangle these possibilities, a single-burst framework was implemented to isolate discrete alpha events and quantify their amplitude, frequency, duration, and abundance. This burst-resolved approach dissociates oscillatory strength from temporal organization, allowing identification of the specific dimensions of alpha activity underlying group differences. As in the time-averaged analyses, it was explicitly tested whether siblings exhibited intermediate values.

### Reduced abundance in autism with an intermediate sibling profile

To determine whether the reduction in alpha power observed with traditional spectral approaches reflected weaker oscillatory amplitude or alterations in the temporal expression of alpha activity, we next characterized the dynamics of discrete alpha events. We quantified alpha bursts using the extended Better OSCillation detection method (eBOSC; (Kosciessa et al., 2020)), an updated implementation of the BOSC framework (Caplan et al., 2001; Whitten et al., 2011). Rather than treating oscillations as continuously present, this method identifies rhythmic episodes in time, allowing separation of how strongly alpha is expressed from how often it occurs.

Within this framework, bursts are defined as periods during which wavelet-derived power at a given frequency exceeds a threshold estimated from the aperiodic (1/f) background (Fig. 2). To ensure that transient fluctuations are not misclassified as oscillations, detections must additionally persist for a minimum duration, here set to at least two cycles at the frequency of interest. By isolating individual events, this approach provides access to complementary metrics of oscillatory organization, including burst abundance, duration, amplitude, and inter-burst intervals.

**Figure 2.**
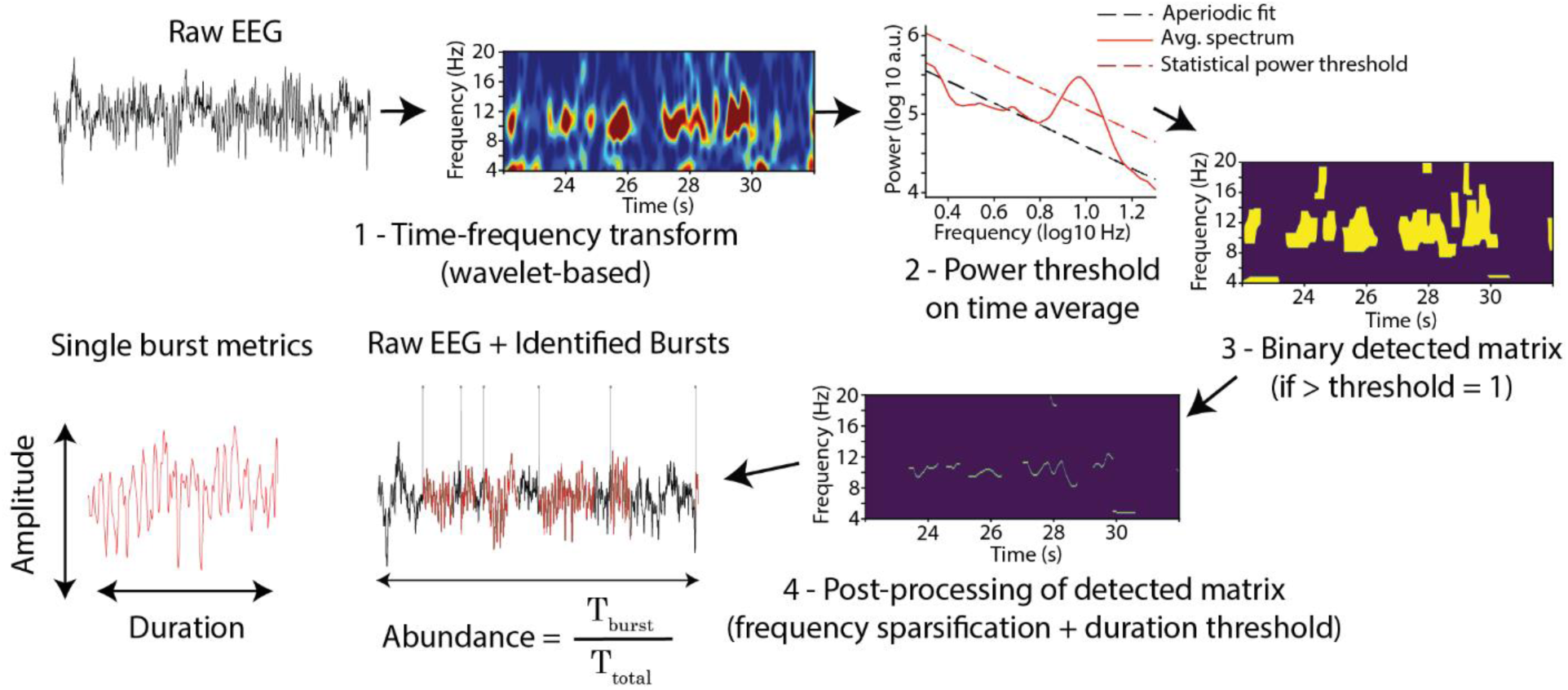
Identification of single alpha events. Overview of the procedure used to detect individual alpha bursts. (1) The continuous EEG signal is transformed into the time–frequency domain. (2) A power spectral density (PSD) estimate is derived, and the aperiodic background is modeled (after excluding the alpha peak) to determine a power threshold. (3) Time-frequency points exceeding this threshold are identified and marked in a binary “detected” matrix. (4) This matrix is then refined by enforcing a minimum duration criterion (here, >2 cycles) and by grouping consecutive points into continuous episodes after determining the local frequency of maximal power for each time point. (5) The resulting onsets and offsets of each episode are projected back onto the raw signal for illustration.

#### Alpha burst amplitude

Average alpha amplitude within detected bursts (Fig. 3A), did not show significant ordinal progression across groups (ß = 1.3 A.U, SE = 75.85, p = 0.98). However, regional effects were robust. Across participants, alpha burst amplitude showed the expected posterior dominance, with significantly higher alpha in the occipital region relative to parietal cortex (β = 1028 A.U, SE = 29.0, p < .001), and significantly lower alpha over frontal regions relative to parietal cortex (β = −105.4 A.U, SE = 26.4, p < .001). Importantly, no significant Group × Region interactions were detected, indicating that this anterior–posterior gradient was comparable across groups. Age was significantly associated with alpha burst amplitude (Supplementary Fig. 2C; ß = −143.8, SE = 40.1, p < 0.001) reflecting a developmental decrease in burst amplitude, with no evidence of moderation by group. Sex was not significantly associated with alpha burst amplitude.

**Figure 3.**
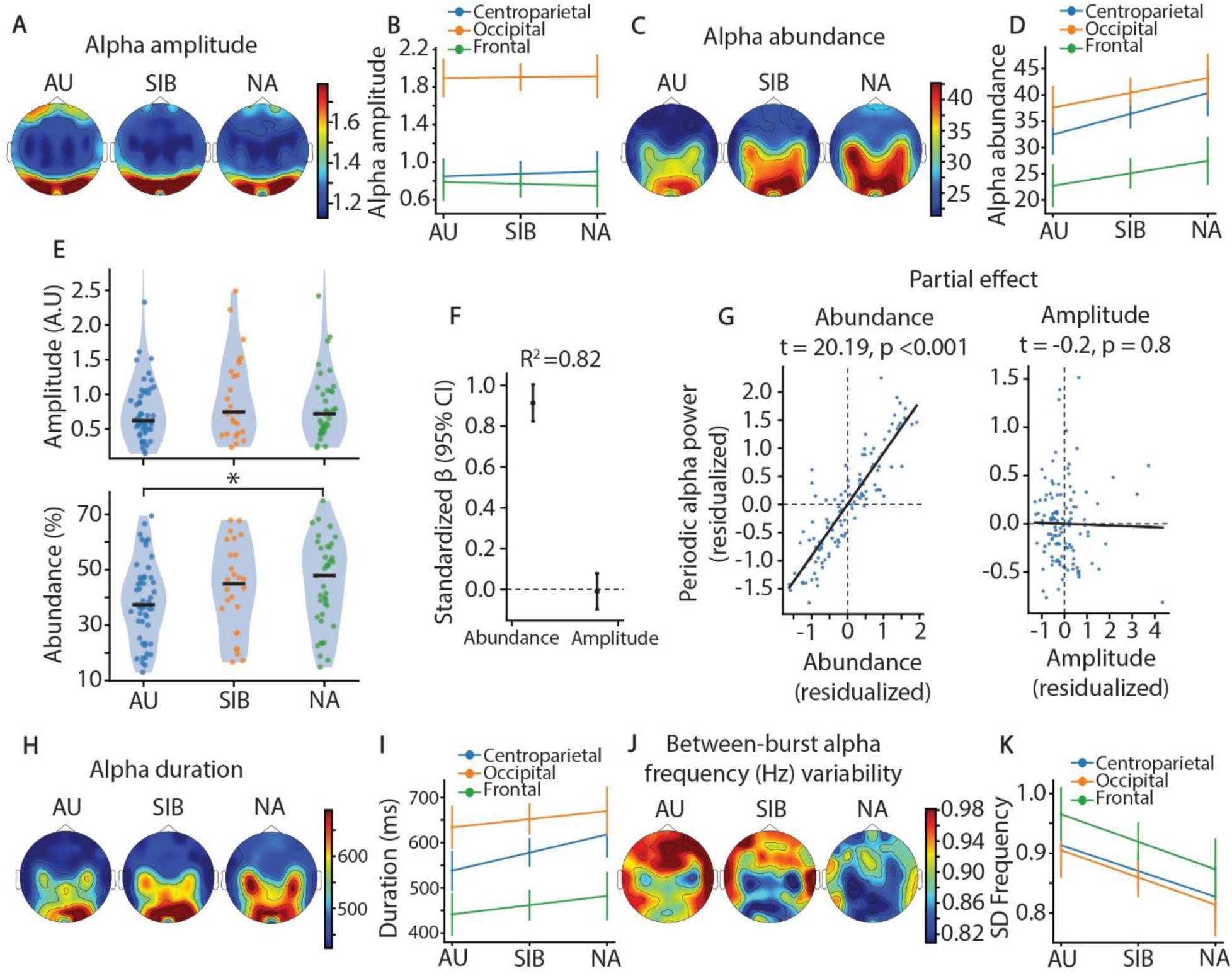
Single-burst analysis of alpha activity reveals reduced alpha abundance at rest in autism leading to a reduction in broadband alpha power. (**A**) Scalp topographies of mean alpha amplitude for autistic (AU; left), sibling (SIB; middle), and non-autistic (NA; right) participants. (**B**) Predicted values from the linear mixed model (LMM) illustrating the absence of ordinal group trend in alpha amplitude. (**C**–**D**) Same conventions as in panels **A** and **B**, but for the abundance of alpha activity (time spent having alpha activity). (**F**) Multiple regression shows the contribution of burst amplitude and burst abundance to periodic alpha power. Standardized coefficients indicate a strong association for abundance (β = 0.91, p < 0.001) and the absence of relationship for amplitude (β < 0.01, p = 0.8). (**G**) Partial regression plots illustrate the unique relationships between periodic alpha power and burst abundance (left) and burst amplitude (right) after accounting for shared variance. (**H**-**I)** and (**J**-**K**) Same conventions as in panels **A** and **B**, but for alpha burst duration and between-burst alpha frequency variability (calculated within participants), respectively.

#### Alpha burst frequency

Across participants, alpha burst frequency exhibited regional variation. Frontal regions showed significantly lower burst frequency relative to parietal cortex (Supplementary Fig. 2D; β = −0.03 Hz, SE = 0.01, p = .002), whereas occipital and parietal regions did not differ significantly (β = 0.019 Hz, SE = 0.01, p = .08). In addition, the right hemisphere was associated with slower alpha burst frequency compared to the left hemisphere (β = −0.027 Hz, SE < 0.01, p < .001). No significant Group trend × Region or Group trend × Hemisphere interactions were observed (Supplementary Fig. 2E), indicating that this spatial pattern was comparable across groups. Age was significantly associated with alpha burst frequency (Supplementary Fig. 2F; β = 0.094 Hz, SE = 0.02, p < .001), reflecting an increase in burst frequency with development. No Age × Group interactions were detected and sex was not significantly associated with alpha burst frequency.

#### Alpha abundance

(defined as the proportion of time occupied by rhythmic alpha episodes) showed a significant positive ordinal trend across groups (β = 4.1%, SE < 0.01, p = .01), indicating increasing abundance from autism → siblings → controls (Fig. 3A, B). Across participants, strong regional effects were observed. Relative to parietal cortex, alpha abundance was significantly reduced over frontal regions (β = −11.3%, SE < 0.01, p < .001) and significantly increased over occipital regions (β = 4.0%, SE < 0.01, p < .001). In addition, the right hemisphere showed slightly higher alpha abundance than the left hemisphere (β = 1.1%, SE < 0.01, p < .001), reflecting a small but highly reliable hemispheric asymmetry. Importantly, the liability pattern was topographically specific. A significant Group trend × Region interaction revealed that the ordinal increase across groups was attenuated at frontal (β = −1.6%, SE < 0.01, p < .001) and occipital electrodes (β = −1.1%, SE < 0.01, p = .01) relative to centroparietal regions. No Group × Hemisphere interaction was observed. Neither age nor sex was significantly associated with alpha abundance.

The categorical model yielded converging results: autistic participants exhibited lower alpha abundance compared to controls (Fig. 3E; β = 8.1%, SE = 0.03, p = .01), whereas siblings did not differ significantly from the autism group (β = 4.0%, SE = 0.3, p = .21), consistent with an intermediate familial profile.

#### Duration and count of alpha burst

Because abundance reflects both how long bursts last and how often they occur, we decompose it into duration and count. Average burst duration (Fig. 3H) showed a significant ordinal effect (Fig. 3I; β = 42ms, SE = 0.01, p = .02), with longer events toward the control end of the continuum. Across participants, strong regional effects were observed. Relative to parietal cortex, duration of alpha burst was significantly shorter for frontal regions (β = −116ms, SE < 0.01, p < .001) and longer for occipital regions (β = 74ms, SE < 0.01, p < .001). Importantly, the liability pattern was topographically specific. A significant Group trend × Region interaction revealed that the ordinal increase across groups was attenuated for both frontal (β = −20ms, SE < 0.01, p = .008) and occipital electrodes (β = −22ms, SE < 0.01, p = .006) relative to centroparietal regions. Neither age nor sex was significantly associated with alpha abundance.

The number of bursts (Supplementary Fig. 2G) followed in the same direction but did not reach conventional significance for the ordinal effect (Supplementary Fig. 2H; β = 1.5, SE = 0.78, p = .054). Across participants, a significant lower number of alpha bursts was found for frontal compared to parietal channels (ß = −4.97, SE = 0.2, p < 0.001) without differences between occipital and parietal regions (ß = 0.08, n.s). No interaction was found between Group trend x Region and Age was not significantly associated with the number of alpha burst.

In the categorical analysis, at centroparietal sites, TD participants exhibited significantly longer burst durations than ASD (β = 83ms, SE = 37, p = .026), whereas siblings did not differ from ASD. Significant Group × Region interactions indicated that spatial gradients differed between groups. In TD, frontal burst durations were significantly shorter relative to parietal cortex than in ASD (β = −38ms, SE = 15, p = .009), indicating a steeper anterior reduction. Conversely, the occipital enhancement relative to parietal cortex was attenuated in TD compared to ASD (β = −47ms, SE = 16, p = .003). No significant interactions were observed for siblings. Age or sex did not contribute to these effects (ß < 5ms, n.s.).

#### Between-burst alpha frequency variability

To further investigate the stability of the alpha generator we analyzed the variability of the amplitude (Supplementary Fig. 2I) and frequency between bursts within participants (Fig. 3J). Alpha frequency variability showed a significant ordinal effect (Fig. 3K; ß = −0.049, SE = 0.01, p = 0.007) indicating increasing variability from autism → siblings → controls. Across participants, the frequency variability was significantly higher for frontal compared to parietal region (β = 0.049 Hz, SE < 0.01, p < .001) with no differences between occipital and parietal regions. Age and sex were not significantly associated with alpha burst variability.

The categorical model yielded converging results: autistic participants exhibited higher alpha frequency variability compared to controls (β = −0.098 Hz, SE = 0.03, p = 0.007), whereas siblings did not differ significantly from the autism group (β = −0.057 Hz, SE = 0.4, p = .17), consistent with an intermediate familial profile.

Between-burst amplitude variability did not show any group effects but showed a negative association with Age with decreasing variability for older participants across groups (Supplementary Fig. 2K, ß = −0.061, SE = 0.1, p = 0.001).

#### Linking burst metrics to broadband alpha power

We then asked which burst characteristics accounted for most of the observed decrease in relative alpha power/periodic alpha power using time-average traditional methods. A multiple regression including burst abundance and burst amplitude as simultaneous predictors explained the vast majority of variance in relative alpha power (Supplementary Fig. 3; R² = .89) and periodic alpha power (Fig. 3F-G). For relative alpha power, although both predictors contributed significantly, the association was markedly stronger for abundance (β = 0.86, t = 23.97) than for amplitude (β = 0.17, t = 4.75). Concerning periodic alpha power, only abundance contributed significantly (Fig. 3F; ß = 0.91, SE = 0.04, p < 0.001; for amplitude: ß < 0.01). Together, these results indicate that time-averaged reduced alpha power in autism (and the intermediate values observed in siblings) primarily reflects spending less time in rhythmic alpha states rather than producing intrinsically weaker oscillations.

Finally, we examined whether the alpha burst metrics that differed between groups were associated with symptom severity within the autism group. For this analysis, alpha parameters were averaged across bilateral centroparietal channels for each participant, and separate linear regressions were conducted within the autistic group only (n = 49). None of the burst-resolved metrics (alpha abundance, average burst duration, or between-burst alpha frequency variability) were significantly associated with SRS-2 total scores (all p_s_ > .80). These findings indicate that reduced alpha engagement in autism does not scale with variation in characteristics associated with autism as assessed by the SRS. Alpha abundance and burst duration were also not significantly associated with FSIQ (both p > .10). A trend-level association was observed between lower FSIQ and greater alpha frequency variability (β = −0.27, SE = 0.14, p = .067), suggesting that increased instability of alpha frequency may relate to cognitive ability, although this effect did not reach statistical significance.

## Discussion

Alpha oscillations dynamically regulate cortical excitability and are widely conceptualized as a neural gating mechanism. Decreases in alpha power are associated with heightened cortical excitability and enhanced sensory processing, whereas increases in alpha reflect stronger inhibitory engagement and reduced sensory responsiveness (Banerjee et al., 2011; Hanslmayr et al., 2011; Kelly et al., 2009; Klimesch et al., 2007; Pfurtscheller & Lopes da Silva, 1999; Snyder & Foxe, 2010). A recurring finding in the resting-state autism literature is a reduction in relative alpha power. However, conventional spectral measures do not clarify whether such reductions reflect weaker oscillatory events or altered temporal organization of alpha activity. In the present study, we addressed this question by decomposing alpha oscillations into burst-resolved features (amplitude, frequency, duration, and rate of occurrence) thereby providing a temporally precise characterization of alpha dynamics beyond traditional time-averaged metrics.

Using conventional spectral approaches, we first replicated the commonly reported reduction in resting-state alpha power in autistic participants (Billeci et al., 2013; Neo et al., 2023; Newson & Thiagarajan, 2018; Wang et al., 2013), observed both in relative alpha power and in periodic alpha power after removal of the aperiodic (1/f) component. Importantly, unaffected siblings exhibited intermediate values, consistent with a liability or endophenotype model. This graded pattern was most pronounced over centroparietal and occipital regions and attenuated over frontal sites, where alpha power was lower across all groups.

Critically, the burst-resolved analyses clarified the source of this reduction. Alpha burst amplitude followed the expected posterior topography (with stronger bursts over occipital channels) but did not differ between groups. In contrast, alpha abundance (the proportion of time spent in an oscillatory alpha state) showed a significant ordinal decrease from controls to siblings to autistic participants, particularly over centroparietal and occipital regions. This reduction in abundance was primarily driven by shorter burst durations, which exhibited a similar graded pattern. To directly test which burst features accounted for reductions in time-averaged alpha power, we conducted regression analyses linking burst metrics to conventional spectral measures. These analyses revealed that alpha abundance, rather than burst amplitude, explained most of the variance in relative and periodic alpha power. In addition, the variability of alpha frequency across bursts was significantly higher in autistic participants and followed a liability pattern, further suggesting reduced stability of alpha-generating mechanisms.

Together, these findings indicate that altered alpha activity in autism reflects reduced temporal stability and persistence of alpha events rather than diminished oscillatory strength per se. In other words, the neural generators of alpha appear capable of producing oscillations of typical amplitude, but the system enters and maintains this inhibitory state less consistently. Reduced persistence of alpha rhythms may therefore index altered regulation of cortical excitability and sensory gating in autism.

### Shorter Alpha Duration as a Potential Mechanism of Cortical Hyper-reactivity

The preservation of burst amplitude alongside reduced burst duration carries potentially important mechanistic implications. Alpha amplitude is often interpreted as indexing the instantaneous strength of inhibitory engagement within a neural ensemble (Klimesch et al., 2007), whereas burst duration reflects how long this inhibitory state is sustained. In this context, our findings raise the possibility that, although inhibitory events can be generated with typical strength, they may be less persistent in autistic children during rest.

This profile aligns with models proposing altered excitatory–inhibitory regulation in autism (Contractor et al., 2021; Takarae & Sweeney, 2017). Such accounts are often discussed in relation to alterations in excitatory–inhibitory balance, potentially involving neurotransmitter systems such as gamma-aminobutyric acid (GABA) and glutamate (McCormick et al., 1993). While findings remain heterogeneous (Ahlfors et al., 2024; Darrell et al., 2025), evidence suggests that atypical GABAergic signaling may contribute to altered regulation of cortical excitability in autism under some circumstances (Pizzarelli & Cherubini, 2011; Robertson et al., 2016). In particular, disruptions in inhibitory circuits, including those involving the thalamic reticular nucleus, have been proposed as one possible mechanism through which sensory gating and cortical excitability could be altered, potentially leading to heightened sensory responsiveness (Mouchati et al., 2019; Ward et al., 2024; Ward et al., 2017).

Within this framework, one possibility is that shorter alpha bursts reflect inhibitory states that are not sustained for as long, increasing the likelihood that cortical circuits operate in higher-gain modes in which sensory input is processed more intensely. If so, this could provide a plausible neural substrate for sensory hyper-reactivity and reduced filtering frequently reported in autism (MacLennan et al., 2022; Taels et al., 2023). However, this interpretation remains speculative and will need to be tested more directly in future work. More generally, our findings raise the possibility that reduced alpha activity in autism may reflect an alteration in temporal organization rather than a simple reduction in oscillatory amplitude (Neo et al., 2023; Wang et al., 2013).

### Convergence with Sleep Spindle Findings

Additional, albeit indirect, support for the idea of reduced inhibitory stability comes from observations of sleep spindle alterations in autism. Sleep spindles are transient thalamocortical oscillations that occur during non-rapid eye movement (NREM) sleep and are thought to play a key role in sensory gating (De Gennaro & Ferrara, 2003). Although spindles occur during NREM sleep and alpha bursts during wakefulness, the two rhythms share critical mechanistic features: both arise from thalamocortical loops, both depend on precise GABAergic timing, and both function as sensory gates, suppressing the propagation of external input (Fernandez & Lüthi, 2020). Notably, sleep studies in autistic children have documented reduced spindle density, shorter spindle duration, and weaker thalamocortical coupling (Farmer et al., 2018; Merikanto et al., 2019; Mylonas et al., 2022; Tessier et al., 2015). Developmental trajectories also parallel one another: both spindles and alpha rhythms show increased frequency and reduced power with age (Mylonas et al., 2022). Moreover, spindle characteristics such as density and duration relate to cognitive abilities, including IQ and adaptive behavior (Farmer et al., 2018), mirroring the associations we observe between alpha dynamics and cognitive measures. Taken together, these converging observations raise the possibility of a broader alteration in state-regulating thalamocortical inhibitory rhythms in autism. In this view, reduced temporal stability of alpha bursts during wakefulness and altered spindle dynamics during sleep could reflect related mechanisms, potentially contributing to both atypical sensory responsivity during wake and the sleep disturbances that are common in autism. However, future work will be needed to directly explore whether waking alpha dynamics and sleep spindle alterations arise from a shared underlying mechanism.

### Developmental Effects and Dimensional Associations

Alpha dynamics showed expected developmental relationships (Hill et al., 2022; Miskovic et al., 2015). We observed age-related decreases in alpha power and increases in alpha frequency across all groups (Supplementary Fig. 1C & Supplementary Fig. 2C). In contrast to some prior reports suggesting attenuated developmental shifts in autism (Edgar et al., 2019; Edgar et al., 2015), we observed preserved age–frequency coupling in autistic, sibling, and non-autistic participants. Further studies with larger and more diverse samples will help refine these conclusions and determine how alpha dynamics may differentially support development across neurodiverse populations.

Although alpha burst abundance, duration, and frequency variability exhibited clear group-level and liability effects, they were not associated with symptom severity within the autism group. These findings suggest that altered alpha burst dynamics reflect a trait-level neurophysiological characteristic rather than a marker of continuous variation in symptom severity.

### Limitations and future directions

Several limitations warrant consideration. First, the functional interpretation of resting-state alpha dynamics remains incomplete. Although reduced alpha abundance might plausibly relate to increased sensory defensiveness or altered sensory regulation, our dataset does not include sensory sensitivity scores or other behavioral measures directly capturing this dimension. More broadly, while burst-resolved measures such as alpha abundance, duration, and amplitude likely reflect meaningful aspects of spontaneous neural organization, their precise significance for cognition and perception during rest is not yet fully understood. Accordingly, the present findings demonstrate group differences in the temporal organization of resting-state alpha activity, but they do not by themselves establish the functional consequences of these differences. Thus, our interpretation in terms of atypical sensory regulation should be viewed as consistent with, but not explanatory of, sensory phenotypes in autism. Future studies combining burst-resolved resting-state analyses with physiological measures (e.g., eye-tracking, ECG, electrodermal activity), validated sensory questionnaires, and task-based indices of sensory processing will be necessary to bridge this mechanistic gap. Second, our autistic sample consisted primarily of cognitively able participants. It is important to determine whether these findings generalize to individuals with higher support needs, for whom differences in neural excitability and sensory processing may be even more pronounced. Third, while the eBOSC approach effectively dissociates periodic from aperiodic activity, it does not fully resolve the spatial origin of alpha bursts. Our resting-state results revealed a clear distinction between occipital and bilateral centroparietal alpha generators, raising the possibility that some effects, particularly those localized to left centroparietal regions, could relate to sensorimotor (mu) activity (Bender et al., 2025). Although differences in movement during rest are unlikely to account for these findings, especially given strong task compliance in cognitively able autistic children, future studies incorporating more refined spatial isolation, such as source-resolved MEG, cortical modeling, or multimodal recordings, will be critical to determine whether altered burst dynamics are specific to particular alpha generators.

Recent findings also offer an interesting contrast to our results. (Murray et al., 2025) reported increased absolute alpha power in autistic adults during both rest and attentional tasks, despite no differences in relative alpha power. This raises the possibility that alpha dynamics evolve across development in autism. Consistent with this view, (Mathewson et al., 2012) observed higher alpha power in autistic adults during eyes-open conditions, with no group differences during eyes-closed rest, suggesting reduced alpha reactivity. Together, these findings point to the possibility that autism is characterized by age-dependent alterations in alpha regulation, with reduced alpha engagement in childhood and a shift toward stronger or more persistent alpha activity in adulthood. Longitudinal and developmental studies using burst-resolved approaches will be critical to determine whether the shorter alpha bursts observed here persist across development or give way to qualitatively different alpha dynamics later in life. Finally, the present findings are based on one minute of eyes-open resting-state data. Although this duration falls within the range commonly used in resting-state studies, it lies at the lower end of typical acquisition lengths. While one minute appears sufficient to capture robust burst activity, longer recordings and larger independent datasets will be needed to establish the stability and generalizability of these findings.

## Conclusion

By decomposing alpha activity into its temporal components, this study reveals a specific alteration in autism: alpha bursts are shorter and less abundant during rest, with siblings exhibiting intermediate profiles. These findings refine current models of neural excitability in autism by indicating that atypical sensory processing may arise from reduced stability of inhibitory states rather than weaker inhibition per se. Burst-resolved analyses therefore provide a powerful framework for identifying mechanistic alterations in cortical excitability and sensory regulation in autism.

## Author contributions

T.V. and S.M. conceived the study. T.V and M.D preprocessed the data and analyzed the data under the supervision of S.M, M.Q and C.B. T.V wrote the first draft of the manuscript. S.M, M.Q and J.J.F edited the manuscript.

## Acknowledgments

We thank Dennis Cregin, Trinca Lecaj, and Daniella Coen for essential contributions to participant recruitment and data acquisition. This work was supported by a grant from the Simons Foundation Autism Research Initiative (SFARI Award # 874845, SM). Support for recruitment and phenotyping of participants was provided by the Human Clinical Phenotyping Core of the NICHD funded Rose. F. Kennedy Intellectual and Developmental Disabilities Research Center (P50 HD105352, SM). JF’s work on this project was supported in part by the Golisano Intellectual and Developmental Disabilities Research Institute (UR-IDDRC), through a center grant from the Eunice Kennedy Shriver National Institute of Child Health and Human Development (P50 HD103536, JJF). The content is solely the responsibility of the authors and does not necessarily represent the official views of the National Institutes of Health.

## Ethics approval and consent to participate

This study was approved by the Institutional Review Board of the Albert Einstein College of Medicine (IRB # 2021-13433). All participants assented to the procedures and parents/guardians provided informed consent.

## Availability of data and materials

The dataset supporting the conclusions of this article is available in the ‘SFARI_EEG_multi-paradigm dataset’ repository (BIDS format), doi:10.18112/openneuro.ds006780.v1.0.0. The scripts used for preprocessing and analyses of the data are available on GitHub: https://github.com/tvanneau/SFARI-Alpha.

---

**Supplementary figure 1.**
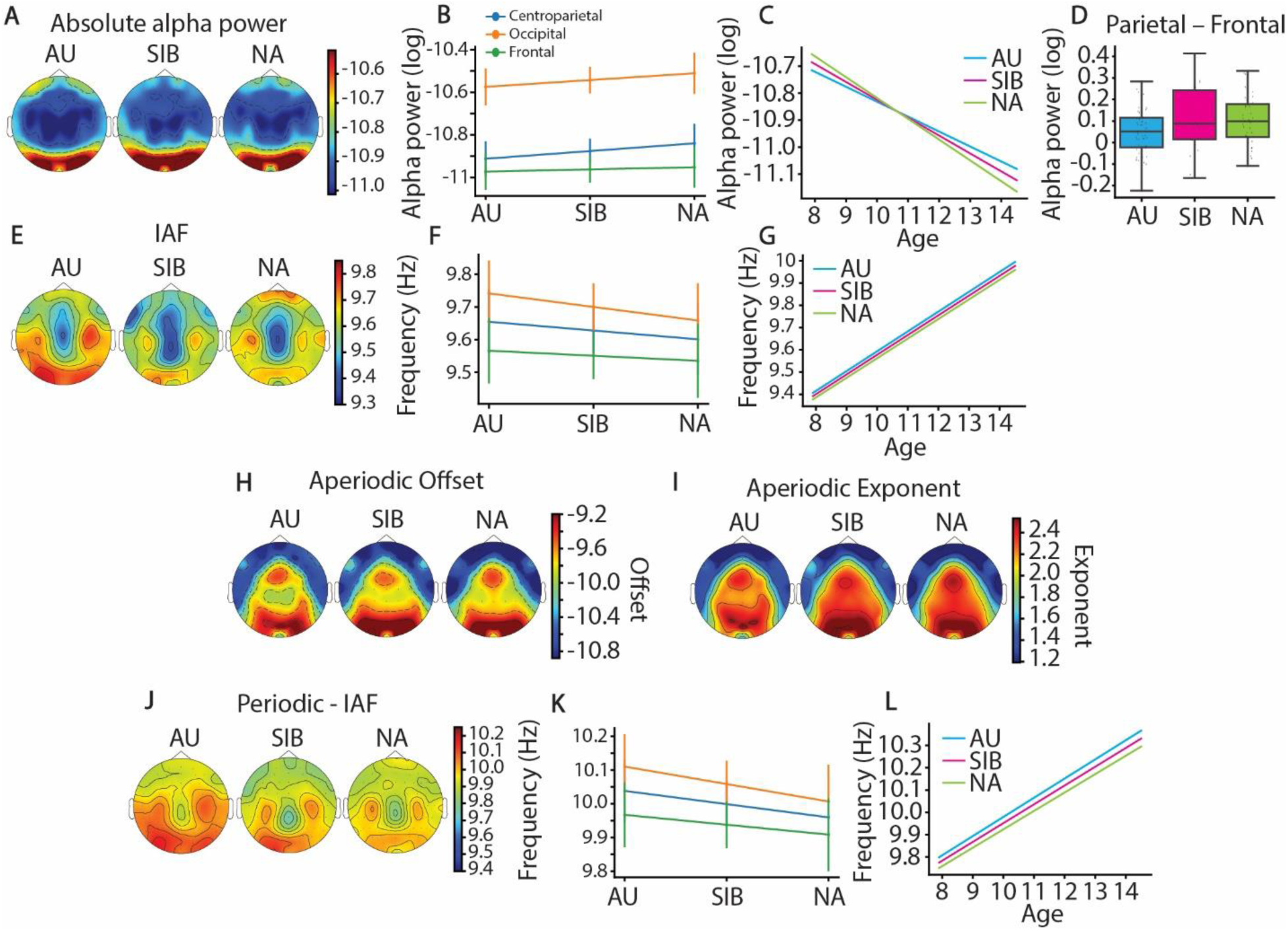
Complementary analysis for band-power and aperiodic/periodic parameters. **(A)** Scalp topographies of absolute alpha power (log-transformed) for autistic participants (AU; left), unaffected siblings (SIB; middle), and non-autistic controls (NA; right). (**B**) Predicted values from the linear mixed model (LMM) illustrating the absence of ordinal group trend in absolute alpha power. (**C**) Predicted values from the LMMs illustrating the effect of age for AU (turquoise), SIB (magenta) and NA (green). (**D**) Boxplots showing the difference between parietal and frontal channels in absolute alpha power for each group. (**E**-**G**) Same analyses as in panels **A**–**C**, respectively, but for individual alpha frequency (IAF). (**H**-**I**) Scalp topographies of the aperiodic offset (**H**) and exponent (**I**). (**J**-**L**) Same analyses as in panels **A**–**C**, respectively, but for IAF estimated after removal of the aperiodic component.

**Supplementary figure 2.**
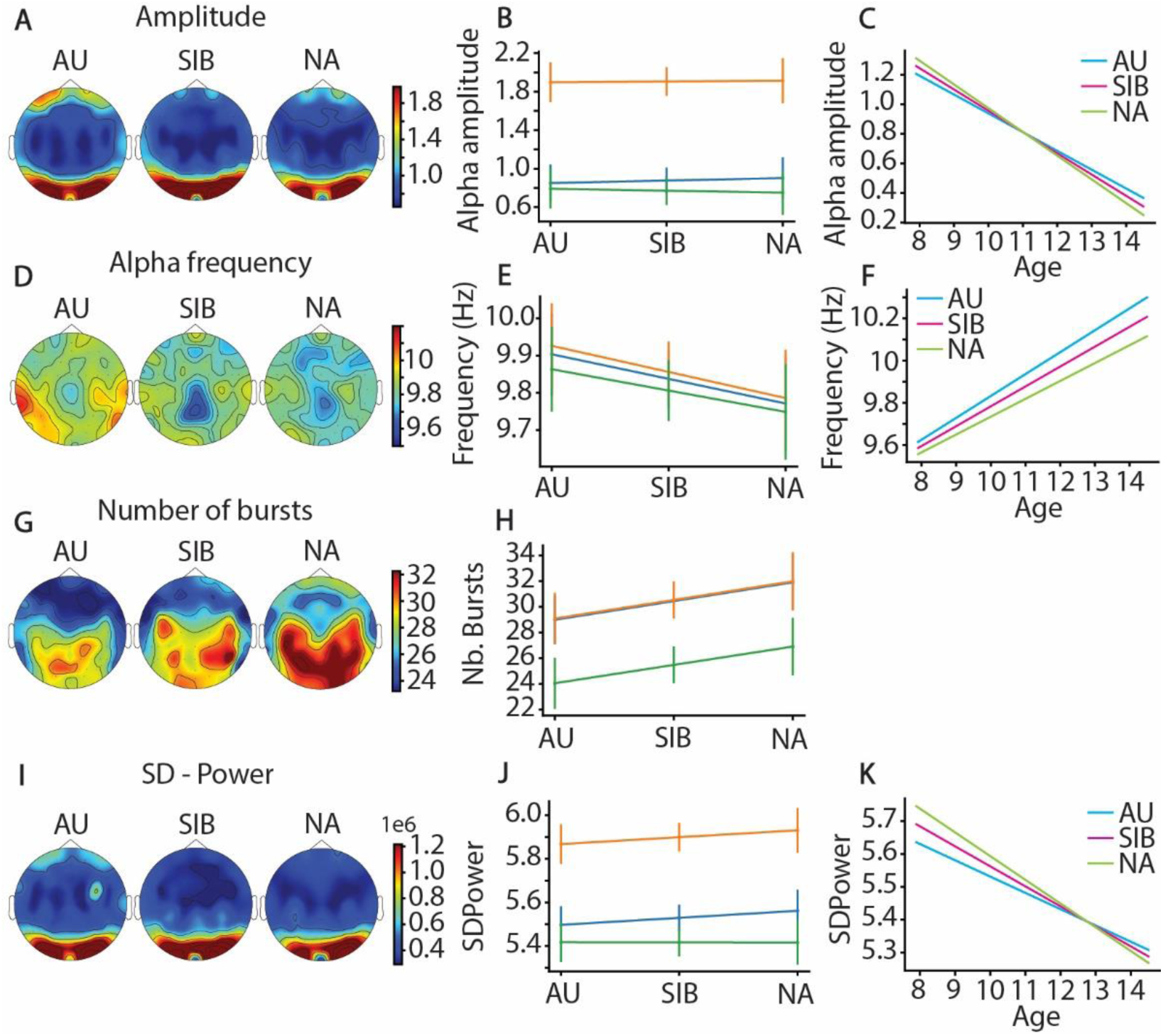
Complementary results for bursts-resolved alpha metrics. (**A**) Scalp topographies of mean alpha amplitude for autistic (AU; left), unaffected siblings (SIB; middle), and non-autistic controls (NA; right) participants. (**B**) Predicted values from the linear mixed model (LMM) illustrating the absence of ordinal group trend in alpha burst amplitude. (**C**) Predicted values from the LMMs illustrating the effect of age for AU (turquoise), SIB (magenta) and NA (green). (**D**–**F**) Same analyses as in panels **A**–**C**, respectively, but for alpha burst frequency. (**G**–**H**) Same analyses as in panels **A**–**B**, respectively, but for the number of alpha bursts. (**I**–**K**) Same analyses as in panels **A**–**C**, respectively, but for the standard deviation of alpha burst amplitude across events within individuals.

**Supplementary figure 3.**
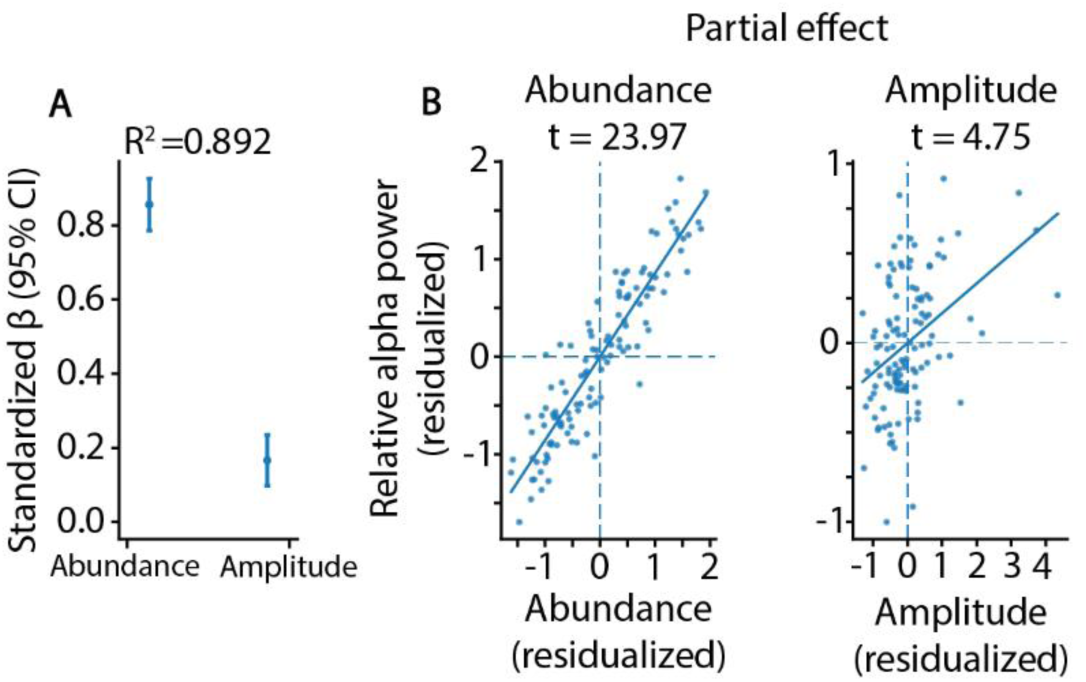
(**A**) Multiple regression showing the contribution of burst abundance and burst amplitude to broadband relative alpha power. Standardized coefficients indicate a strong association for abundance (β = 0.86) and a weaker but significant contribution for amplitude (β = 0.17; both p < .001; R² = .89). (**B**) Partial regression plots illustrating the unique relationships between relative alpha power and burst abundance (left) and burst amplitude (right) after accounting for shared variance.

## Footnotes

1 In this paper, we primarily adopt identity-first language, reflecting prevailing preferences among many autistic adults and self-advocates. We recognize, however, that person-first language is preferred by others in the autism community.

## References

Ahlfors, S. P., Graham, S., Bharadwaj, H., Mamashli, F., Khan, S., Joseph, R. M., Losh, A., Pawlyszyn, S., McGuiggan, N. M., Vangel, M., Hamalainen, M. S., & Kenet, T. (2024). No Differences in Auditory Steady-State Responses in Children with Autism Spectrum Disorder and Typically Developing Children. J Autism Dev Disord, 54(5), 1947–1960. 10.1007/s10803-023-05907-w

American Psychiatric, A. (2022). Diagnostic and Statistical Manual of Mental Disorders. 10.1176/appi.books.9780890425787

Banerjee, S., Snyder, A. C., Molholm, S., & Foxe, J. J. (2011). Oscillatory alpha-band mechanisms and the deployment of spatial attention to anticipated auditory and visual target locations: supramodal or sensory-specific control mechanisms? J Neurosci, 31(27), 9923–9932. 10.1523/JNEUROSCI.4660-10.2011

Belmonte, M. K. (2017). Obligatory Processing of Task-Irrelevant Stimuli: A Hallmark of Autistic Cognitive Style Within and Beyond the Diagnosis. Biol Psychiatry Cogn Neurosci Neuroimaging, 2(6), 461–463. 10.1016/j.bpsc.2017.07.002

Bender, A., Voytek, B., & Schaworonkow, N. (2025). Resting-state Alpha and Mu Rhythms Change Shape across Development But Lack Diagnostic Sensitivity for Attention-Deficit/Hyperactivity Disorder and Autism. J Cogn Neurosci, 1–35. 10.1162/jocn_a_02323

Bigdely-Shamlo, N., Mullen, T., Kothe, C., Su, K. M., & Robbins, K. A. (2015). The PREP pipeline: Standardized preprocessing for large-scale EEG analysis. Frontiers in Neuroinformatics, 9(JUNE), 1–19. 10.3389/fninf.2015.00016

Billeci, L., Sicca, F., Maharatna, K., Apicella, F., Narzisi, A., Campatelli, G., Calderoni, S., Pioggia, G., & Muratori, F. (2013). On the application of quantitative EEG for characterizing autistic brain: a systematic review. Front Hum Neurosci, 7, 442. 10.3389/fnhum.2013.00442

Caplan, J. B., Madsen, J. R., Raghavachari, S., & Kahana, M. J. (2001). Distinct patterns of brain oscillations underlie two basic parameters of human maze learning. J Neurophysiol, 86(1), 368–380. 10.1152/jn.2001.86.1.368

Carter Leno, V., Tomlinson, S. B., Chang, S. A., Naples, A. J., & McPartland, J. C. (2018). Resting-state alpha power is selectively associated with autistic traits reflecting behavioral rigidity. Sci Rep, 8(1), 11982. 10.1038/s41598-018-30445-2

Contractor, A., Ethell, I. M., & Portera-Cailliau, C. (2021). Cortical interneurons in autism. Nat Neurosci, 24(12), 1648–1659. 10.1038/s41593-021-00967-6

Darrell, M., Vanneau, T., Cregin, D., Lecaj, T., Foxe, J. J., & Molholm, S. (2025). Testing the auditory steady-state response (ASSR) to 40-Hz and 27-Hz click trains in children with autism spectrum disorder and their first-degree biological relatives: A high-density electroencephalographic (EEG) study. bioRxiv. 10.1101/2025.08.05.668742

Dawson, G., Toth, K., Abbott, R., Osterling, J., Munson, J., Estes, A., & Liaw, J. (2004). Early social attention impairments in autism: social orienting, joint attention, and attention to distress. Dev Psychol, 40(2), 271–283. 10.1037/0012-1649.40.2.271

Dawson, G., Webb, S. J., Carver, L., Panagiotides, H., & McPartland, J. (2004). Young children with autism show atypical brain responses to fearful versus neutral facial expressions of emotion. Dev Sci, 7(3), 340–359. 10.1111/j.1467-7687.2004.00352.x

De Gennaro, L., & Ferrara, M. (2003). Sleep spindles: an overview. Sleep Med Rev, 7(5), 423–440. 10.1053/smrv.2002.0252

Dockree, P. M., Kelly, S. P., Foxe, J. J., Reilly, R. B., & Robertson, I. H. (2007). Optimal sustained attention is linked to the spectral content of background EEG activity: greater ongoing tonic alpha (approximately 10 Hz) power supports successful phasic goal activation. Eur J Neurosci, 25(3), 900–907. 10.1111/j.1460-9568.2007.05324.x

Donoghue, T., Haller, M., Peterson, E. J., Varma, P., Sebastian, P., Gao, R., Noto, T., Lara, A. H., Wallis, J. D., Knight, R. T., Shestyuk, A., & Voytek, B. (2020). Parameterizing neural power spectra into periodic and aperiodic components. Nat Neurosci, 23(12), 1655–1665. 10.1038/s41593-020-00744-x

Donoghue, T., Schaworonkow, N., & Voytek, B. (2022). Methodological considerations for studying neural oscillations. European Journal of Neuroscience, 55(11-12), 3502–3527. 10.1111/ejn.15361

Edgar, J. C., Dipiero, M., McBride, E., Green, H. L., Berman, J., Ku, M., Liu, S., Blaskey, L., Kuschner, E., Airey, M., Ross, J. L., Bloy, L., Kim, M., Koppers, S., Gaetz, W., Schultz, R. T., & Roberts, T. P. L. (2019). Abnormal maturation of the resting-state peak alpha frequency in children with autism spectrum disorder. Human Brain Mapping, 40(11), 3288–3298. 10.1002/hbm.24598

Edgar, J. C., Heiken, K., Chen, Y. H., Herrington, J. D., Chow, V., Liu, S., Bloy, L., Huang, M., Pandey, J., Cannon, K. M., Qasmieh, S., Levy, S. E., Schultz, R. T., & Roberts, T. P. L. (2015). Resting-State Alpha in Autism Spectrum Disorder and Alpha Associations with Thalamic Volume. Journal of Autism and Developmental Disorders, 45(3), 795–804. 10.1007/s10803-014-2236-1

Farmer, C. A., Chilakamarri, P., Thurm, A. E., Swedo, S. E., Holmes, G. L., & Buckley, A. W. (2018). Spindle activity in young children with autism, developmental delay, or typical development. Neurology, 91(2), e112–e122. 10.1212/WNL.0000000000005759

Feingold, J., Gibson, D. J., DePasquale, B., & Graybiel, A. M. (2015). Bursts of beta oscillation differentiate postperformance activity in the striatum and motor cortex of monkeys performing movement tasks. Proc Natl Acad Sci U S A, 112(44), 13687–13692. 10.1073/pnas.1517629112

Fernandez, L. M. J., & Lüthi, A. (2020). Sleep Spindles: Mechanisms and Functions. Physiological Reviews, 100(2), 805–868. 10.1152/physrev.00042.2018

Foxe, J. J., Simpson, G. V., & Ahlfors, S. P. (1998). Parieto-occipital approximately 10 Hz activity reflects anticipatory state of visual attention mechanisms. NeuroReport, 9(17), 3929–3933. 10.1097/00001756-199812010-00030

Foxe, J. J., & Snyder, A. C. (2011). The role of alpha-band brain oscillations as a sensory suppression mechanism during selective attention. Frontiers in Psychology, 2(JUL). 10.3389/fpsyg.2011.00154

Gramfort, A., Luessi, M., Larson, E., Engemann, D. A., Strohmeier, D., Brodbeck, C., Goj, R., Jas, M., Brooks, T., Parkkonen, L., & Hamalainen, M. (2013). MEG and EEG data analysis with MNE-Python. Front Neurosci, 7, 267. 10.3389/fnins.2013.00267

Haegens, S., Nacher, V., Luna, R., Romo, R., & Jensen, O. (2011). alpha-Oscillations in the monkey sensorimotor network influence discrimination performance by rhythmical inhibition of neuronal spiking. Proc Natl Acad Sci U S A, 108(48), 19377–19382. 10.1073/pnas.1117190108

Hanslmayr, S., Gross, J., Klimesch, W., & Shapiro, K. L. (2011). The role of alpha oscillations in temporal attention. Brain Res Rev, 67(1-2), 331–343. 10.1016/j.brainresrev.2011.04.002

Hill, A. T., Clark, G. M., Bigelow, F. J., Lum, J. A. G., & Enticott, P. G. (2022). Periodic and aperiodic neural activity displays age-dependent changes across early-to-middle childhood. Dev Cogn Neurosci, 54, 101076. 10.1016/j.dcn.2022.101076

Iemi, L., Gwilliams, L., Samaha, J., Auksztulewicz, R., Cycowicz, Y. M., King, J. R., Nikulin, V. V., Thesen, T., Doyle, W., Devinsky, O., Schroeder, C. E., Melloni, L., & Haegens, S. (2022). Ongoing neural oscillations influence behavior and sensory representations by suppressing neuronal excitability. NeuroImage, 247. 10.1016/j.neuroimage.2021.118746

Jensen, O., & Mazaheri, A. (2010). Shaping functional architecture by oscillatory alpha activity: gating by inhibition. Front Hum Neurosci, 4, 186. 10.3389/fnhum.2010.00186

Jones, S. R. (2016). When brain rhythms aren’t ‘rhythmic’: implication for their mechanisms and meaning. Curr Opin Neurobiol, 40, 72–80. 10.1016/j.conb.2016.06.010

Keehn, B., Muller, R. A., & Townsend, J. (2013). Atypical attentional networks and the emergence of autism. Neurosci Biobehav Rev, 37(2), 164–183. 10.1016/j.neubiorev.2012.11.014

Keehn, B., Nair, A., Lincoln, A. J., Townsend, J., & Muller, R. A. (2016). Under-reactive but easily distracted: An fMRI investigation of attentional capture in autism spectrum disorder. Dev Cogn Neurosci, 17, 46–56. 10.1016/j.dcn.2015.12.002

Keehn, B., Westerfield, M., Müller, R. A., & Townsend, J. (2017). Autism, Attention, and Alpha Oscillations: An Electrophysiological Study of Attentional Capture. Biological Psychiatry: Cognitive Neuroscience and Neuroimaging, 2(6), 528–536. 10.1016/j.bpsc.2017.06.006

Kelly, S. P., Gomez-Ramirez, M., & Foxe, J. J. (2009). The strength of anticipatory spatial biasing predicts target discrimination at attended locations: a high-density EEG study. Eur J Neurosci, 30(11), 2224–2234. 10.1111/j.1460-9568.2009.06980.x

Klimesch, W. (1999). EEG alpha and theta oscillations reflect cognitive and memory performance: a review and analysis. Brain Res Brain Res Rev, 29(2-3), 169–195. 10.1016/s0165-0173(98)00056-3

Klimesch, W., Sauseng, P., & Hanslmayr, S. (2007). EEG alpha oscillations: The inhibition-timing hypothesis. In Brain Research Reviews (Vol. 53, pp. 63–88).

Kosciessa, J. Q., Grandy, T. H., Garrett, D. D., & Werkle-Bergner, M. (2020). Single-trial characterization of neural rhythms: Potential and challenges. NeuroImage, 206, 116331. 10.1016/j.neuroimage.2019.116331

Lord, C., Rutter, M., & Le Couteur, A. (1994). Autism Diagnostic Interview-Revised: a revised version of a diagnostic interview for caregivers of individuals with possible pervasive developmental disorders. J Autism Dev Disord, 24(5), 659–685. 10.1007/BF02172145

Lundqvist, M., Herman, P., Warden, M. R., Brincat, S. L., & Miller, E. K. (2018). Gamma and beta bursts during working memory readout suggest roles in its volitional control. Nat Commun, 9(1), 394. 10.1038/s41467-017-02791-8

Lundqvist, M., Miller, E. K., Nordmark, J., Liljefors, J., & Herman, P. (2024). Beta: bursts of cognition. Trends Cogn Sci, 28(7), 662–676. 10.1016/j.tics.2024.03.010

MacLennan, K., O’Brien, S., & Tavassoli, T. (2022). In Our Own Words: The Complex Sensory Experiences of Autistic Adults. J Autism Dev Disord, 52(7), 3061–3075. 10.1007/s10803-021-05186-3

Mathewson, K. E., Lleras, A., Beck, D. M., Fabiani, M., Ro, T., & Gratton, G. (2011). Pulsed out of awareness: EEG alpha oscillations represent a pulsed-inhibition of ongoing cortical processing. Front Psychol, 2, 99. 10.3389/fpsyg.2011.00099

Mathewson, K. J., Jetha, M. K., Drmic, I. E., Bryson, S. E., Goldberg, J. O., & Schmidt, L. A. (2012). Regional EEG alpha power, coherence, and behavioral symptomatology in autism spectrum disorder. Clinical Neurophysiology, 123(9), 1798–1809. 10.1016/j.clinph.2012.02.061

McCormick, D. A., Wang, Z., & Huguenard, J. (1993). Neurotransmitter control of neocortical neuronal activity and excitability. Cereb Cortex, 3(5), 387–398. 10.1093/cercor/3.5.387

Merikanto, I., Kuula, L., Makkonen, T., Salmela, L., Raikkonen, K., & Pesonen, A. K. (2019). Autistic Traits Are Associated With Decreased Activity of Fast Sleep Spindles During Adolescence. J Clin Sleep Med, 15(3), 401–407. 10.5664/jcsm.7662

Miskovic, V., Ma, X., Chou, C. A., Fan, M., Owens, M., Sayama, H., & Gibb, B. E. (2015). Developmental changes in spontaneous electrocortical activity and network organization from early to late childhood. NeuroImage, 118, 237–247. 10.1016/j.neuroimage.2015.06.013

Mouchati, P. R., Barry, J. M., & Holmes, G. L. (2019). Functional brain connectivity in a rodent seizure model of autistic-like behavior. Epilepsy Behav, 95, 87–94. 10.1016/j.yebeh.2019.03.046

Murphy, J. W., Foxe, J. J., Peters, J. B., & Molholm, S. (2014). Susceptibility to distraction in autism spectrum disorder: Probing the integrity of oscillatory Alpha-Band Suppression Mechanisms. Autism Research, 7(4), 442–458. 10.1002/aur.1374

Murray, S. O., Seczon, D. L., Pettet, M., Rea, H. M., Woodard, K. M., Kolodny, T., & Webb, S. J. (2025). Increased alpha power in autistic adults: Relation to sensory behaviors and cortical volume. Autism Res, 18(1), 56–69. 10.1002/aur.3266

Mylonas, D., Machado, S., Larson, O., Patel, R., Cox, R., Vangel, M., Maski, K., Stickgold, R., & Manoach, D. S. (2022). Dyscoordination of non-rapid eye movement sleep oscillations in autism spectrum disorder. Sleep, 45(3). 10.1093/sleep/zsac010

Neo, W. S., Foti, D., Keehn, B., & Kelleher, B. (2023). Resting-state EEG power differences in autism spectrum disorder: a systematic review and meta-analysis. In Translational Psychiatry (Vol. 13): Springer Nature.

Newson, J. J., & Thiagarajan, T. C. (2018). EEG Frequency Bands in Psychiatric Disorders: A Review of Resting State Studies. Front Hum Neurosci, 12, 521. 10.3389/fnhum.2018.00521

Perrin, Pernier, Bertrand, & Echallier. (1989). Spherical splines for scalp potential and current density mapping. Electroencephalography and Clinical Neurophysiology, 72, 184–187.

Pfurtscheller, G., & Lopes da Silva, F. H. (1999). Event-related EEG/MEG synchronization and desynchronization: basic principles. Clin Neurophysiol, 110(11), 1842–1857. 10.1016/s1388-2457(99)00141-8

Pizzarelli, R., & Cherubini, E. (2011). Alterations of GABAergic Signaling in Autism Spectrum Disorders. Neural Plasticity, 2011, 1–12. 10.1155/2011/297153

Robertson, C. E., & Baron-Cohen, S. (2017). Sensory perception in autism. Nat Rev Neurosci, 18(11), 671–684. 10.1038/nrn.2017.112

Robertson, C. E., Ratai, E. M., & Kanwisher, N. (2016). Reduced GABAergic Action in the Autistic Brain. Curr Biol, 26(1), 80–85. 10.1016/j.cub.2015.11.019

Rutter, M., Le Couteur, A., & Lord, C. (2003). Autism diagnostic interview-revised. Los Angeles, CA: Western Psychological Services, 29(2003), 30.

Sadaghiani, S., & Kleinschmidt, A. (2016). Brain Networks and alpha-Oscillations: Structural and Functional Foundations of Cognitive Control. Trends Cogn Sci, 20(11), 805–817. 10.1016/j.tics.2016.09.004

Sauseng, P., Klimesch, W., Stadler, W., Schabus, M., Doppelmayr, M., Hanslmayr, S., Gruber, W. R., & Birbaumer, N. (2005). A shift of visual spatial attention is selectively associated with human EEG alpha activity. Eur J Neurosci, 22(11), 2917–2926. 10.1111/j.1460-9568.2005.04482.x

Sherman, M. A., Lee, S., Law, R., Haegens, S., Thorn, C. A., Hamalainen, M. S., Moore, C. I., & Jones, S. R. (2016). Neural mechanisms of transient neocortical beta rhythms: Converging evidence from humans, computational modeling, monkeys, and mice. Proc Natl Acad Sci U S A, 113(33), E4885–4894. 10.1073/pnas.1604135113

Snyder, A. C., & Foxe, J. J. (2010). Anticipatory attentional suppression of visual features indexed by oscillatory alpha-band power increases: a high-density electrical mapping study. J Neurosci, 30(11), 4024–4032. 10.1523/JNEUROSCI.5684-09.2010

Steriade, M., Gloor, P., Llinas, R. R., Lopes de Silva, F. H., & Mesulam, M. M. (1990). Report of IFCN Committee on Basic Mechanisms. Basic mechanisms of cerebral rhythmic activities. Electroencephalogr Clin Neurophysiol, 76(6), 481–508. 10.1016/0013-4694(90)90001-z

Taels, L., Feyaerts, J., Lizon, M., De Smet, M., & Vanheule, S. (2023). ’I felt like my senses were under attack’: An interpretative phenomenological analysis of experiences of hypersensitivity in autistic individuals. Autism, 27(8), 2269–2280. 10.1177/13623613231158182

Takarae, Y., & Sweeney, J. (2017). Neural Hyperexcitability in Autism Spectrum Disorders. Brain Sci, 7(10). 10.3390/brainsci7100129

Tessier, S., Lambert, A., Chicoine, M., Scherzer, P., Soulieres, I., & Godbout, R. (2015). Intelligence measures and stage 2 sleep in typically-developing and autistic children. Int J Psychophysiol, 97(1), 58–65. 10.1016/j.ijpsycho.2015.05.003

Van Diepen, R. M., Foxe, J. J., & Mazaheri, A. (2019). The functional role of alpha-band activity in attentional processing: the current zeitgeist and future outlook. Curr Opin Psychol, 29, 229–238. 10.1016/j.copsyc.2019.03.015

van Ede, F., Quinn, A. J., Woolrich, M. W., & Nobre, A. C. (2018). Neural Oscillations: Sustained Rhythms or Transient Burst-Events? Trends Neurosci, 41(7), 415–417. 10.1016/j.tins.2018.04.004

Vanneau, T., Foxe, J. J., Beker, S., & Molholm, S. (2025). Disrupted Top-Down Modulation as a Mechanism of Impaired Multisensory Processing in Children with an Autism Spectrum Diagnosis. bioRxiv. 10.1101/2025.11.13.688243

Vidaurre, D., Quinn, A. J., Baker, A. P., Dupret, D., Tejero-Cantero, A., & Woolrich, M. W. (2016). Spectrally resolved fast transient brain states in electrophysiological data. NeuroImage, 126, 81–95. 10.1016/j.neuroimage.2015.11.047

Wang, J., Barstein, J., Ethridge, L. E., Mosconi, M. W., Takarae, Y., & Sweeney, J. A. (2013). Resting state EEG abnormalities in autism spectrum disorders. Journal of Neurodevelopmental Disorders, 5(1). 10.1186/1866-1955-5-24

Ward, C., Nasrallah, K., Tran, D., Sabri, E., Vazquez, A., Sjulson, L., Castillo, P. E., & Batista-Brito, R. (2024). Developmental disruption of Mef2c in Medial Ganglionic Eminence-derived cortical inhibitory interneurons impairs cellular and circuit function. bioRxiv. 10.1101/2024.05.01.592084

Ward, J., Hoadley, C., Hughes, J. E., Smith, P., Allison, C., Baron-Cohen, S., & Simner, J. (2017). Atypical sensory sensitivity as a shared feature between synaesthesia and autism. Sci Rep, 7, 41155. 10.1038/srep41155

Whitten, T. A., Hughes, A. M., Dickson, C. T., & Caplan, J. B. (2011). A better oscillation detection method robustly extracts EEG rhythms across brain state changes: the human alpha rhythm as a test case. NeuroImage, 54(2), 860–874. 10.1016/j.neuroimage.2010.08.064

Worden, M. S., Foxe, J. J., Wang, N., & Simpson, G. V. (2000). Anticipatory Biasing of Visuospatial Attention Indexed by Retinotopically Specific-Band Electroencephalography Increases over Occipital Cortex. http://www.jneurosci.org/cgi/content/full/4016

